# Peasants at the queen’s table?: The microbiome’s dynamics throughout swarming preparation in honey bees (*Apis mellifera*)

**DOI:** 10.1101/2024.09.22.614346

**Authors:** Michał. R. Kolasa, Bartłomiej Molasy, Aneta Strachecka, Anna Michalik

## Abstract

The intricate relationship between hosts and bacterial symbionts was crucial for the evolution of multiple insect clades. Western honey bees *(Apis mellifera)* have distinctive and highly stable microbiomes in terms of bacterial taxonomic composition. However, despite the significant development of molecular techniques observed in recent decades, we still are far from understanding much about the dynamics of honey bees’ microbiomes in terms of their strain diversity and quantities. The overall stable composition of their microbiome and complex behavior make honey bees a perfect model species for studying the gut-brain axis, where the gut microbiome can efficiently influence the host’s behavior. Here, by implementing high-throughput amplicon sequencing of bacterial 16S rRNA V4 and V1V2 hypervariable regions alongside a cutting-edge quantification approach, we aimed to describe the dynamic in honey bee young workers’ microbiome composition and quantity throughout swarming preparation.

Our results show no changes in microbial absolute abundances throughout the swarming preparation among young worker bees. The V4 and V1V2 datasets congruently reconstructed microbial composition with some notable exceptions, and differential abundance analysis indicated that *Bombella* and *Bartonella* significantly changed over batches.

## Introduction

Symbiosis with microorganisms was crucial for the evolution on Earth, particularly in the divergence of insects, allowing them to colonize all land habitats and occupy all the trophic niches (Cornwallis et al. 2023). Symbiotic relationships between bacteria and their hosts can be classified on various levels, depending either on the stability of host-symbiont associations (Perreau and Moran 2022) or the nature of the relationship (beneficial, neutral, or harmful) (Hammer et al. 2019). Many insect symbioses are long-established, dating back millions of years ago, and played a key role in their evolution; on the other hand, we have many examples of relatively new symbiont acquisitions. However, there is no doubt that both ancestral and newly acquired symbionts are still important players in shaping insect biology or their adaptation to changing environmental pressure and that this is an ongoing process (Łukasik and Kolasa 2024). The implementation of sequencing techniques in symbioses studies enabled a better understanding of many aspects of their functioning. However, despite these advances, our knowledge of microbiota’s role is often limited to a few model species. One of the best-studied insect species regarding microbiome composition and role is the western honey bee (*Apis mellifera*). Their gut microbiota has been described as highly stable, consisting of only nine taxa (Engel and Moran 2013), with most of the composition patterns and functions being performed by only five core species (Motta and Moran 2024): *Snodgrassella alvi, Gilliamella apicola, Bombilactobacillus* (previously called *Lactobacillus* Firm-4 (Zheng et al. 2020), *Lactobacillus* (previously called *Lactobacillus* Firm-5), and *Bifidobacterium asteroides*. However, variations in workers’ microbiomes have been reported depending on the season (Ludvigsen et al. 2015; Kešnerová et al. 2020), diet (Jones et al. 2018a), or geography (Ludvigsen et al. 2017). Recent studies also highlight the role of specific symbiont strains in nutrition provisioning and pathogen resistance (Miller et al. 2021; Parish et al. 2022). However, the honey bee symbionts’ role is not limited only to triggering immune responses or coping with environmental stressors.

Recent studies suggest that gut microbiota can influence various aspects of honey bees’ behavior. Zhang et al. (2022a) experimentally showed how *Lactobacillus* modulates learning and memory behavior by regulating tryptophan metabolism. Another study (Zhang et al. 2022b) showed how different bacteria taxa regulate specific modules of metabolites in workers’ hemolymph. For example, *Bombilactobacillus* and *Lactobacillus* (previously described as *Lactobacillus* Frim-4 and Firm-5) altered amino acid metabolism pathways, leading to upregulations of genes responsible for olfactory functions and division of labour. At the same time, *Gilliamella* were mainly responsible for circulating metabolites involved in carbohydrate and glycerophospholipid metabolism pathways. Another example is an experiment consisting of mono-colonization with *Bifidobacterium asteroides*, which showed that worker bees colonized with this bacterium had elevated gut concentrations of juvenile hormone III derivatives (Kešnerová et al. 2017). This hormone is responsible for insect growth, development, and reproduction, and in honey bees is a key factor for transitioning from nurse bees to foraging ones. However, the effects of symbionts are not limited to single individuals. Liberti et al. (2022) compared microbiota-colonized and microbiota-deprived bees regarding social behavior. They found that colonized bees have increased levels of brain metabolites (such as ornithine and serine) and increased specialized head-to-head interactions between nestmates. Their complex behavior system and relatively stable microbiota make honey bees an excellent and promising system for studying the gut-brain axis (Liberti and Engel 2020).

Due to their social nature, honey bees have become a model species in behavioral studies. The most widely known phenomenon is their ability to communicate the location of food sources by dancing (Von Frisch 1967). Karl Von Frisch described this phenomenon as “the most astounding example of non-primate communication that we know”, and his discovery secured him a Nobel prize in 1973. After 50 years, our understanding of the bees’ cognitive capabilities (e.g., Seeley et al. 1991; Jaumann et al. 2013; Avarguès-Weber and Giurfa 2014), division of labour (Page and Robinson 1991) has increased significantly. One phenomenon remains poorly understood despite this progress - reproductive swarming (hereafter swarming). This complex and multi-step process leads to the division of the colony, with an old queen leaving the hive with approximately 75% of bees (Fefferman and Starks 2006) and a new queen emerging. Typically, it occurs in mid-spring when the food source is most abundant. Although scientific literature points out mass queen rearing as the first visible sign of swarming preparation (Grozinger et al. 2014), beekeepers mention other signs that inform them about upcoming swarming, such as the emergence of drones and the cessation of building wax combs. Swarming preparations require coordinated behavior between thousands of individuals. Studies suggest that there is no nepotistic behavior during swarming (Châline et al. 2005; Rangel et al. 2009). Therefore, an environmental factor is probably responsible for triggering such preparations. Many factors, such as increased colony size, changes in the age proportion in the colony toward younger bees, limitations of available brood cells, and reduced transmission of queen pheromones, have been hypothesized to push the colony toward preparations to swarm (Winston 1991). However, Grozinger et al. (2014) pointed out that these factors are probably more synergistic, as none alone triggers swarming behavior. Indeed, a modeling approach (Fefferman and Starks 2006) proved that three factors (colony/population size, brood nest congestion, and skewed worker age distribution) generated swarming patterns similar to those obtained in empirical studies. Nonetheless, each model only resulted in swarming after other variables defined by their individual models reached their own thresholds. These results may suggest that an undefined critical cue still triggers swarming and correlates with all the mentioned factors.

Royal jelly is a secretion of nurse worker bees’ hypopharyngeal and mandibular glands (Hassanyar et al. 2023) and is best known for the cast determination in honey bees. However, there is still ongoing scientific debate about which compound of royal jelly is responsible for complex physiological and behavioral changes that distinguish workers and queen bees (Maleszka 2018). Nonetheless, numerous studies proved that royal jelly could change metabolic flux (Foret et al. 2012), hormone levels (Hartfelder and Engels 1998) or launch an epigenetic cascade leading to alteration in DNA methylation, histone modifications, and non-protein coding RNAs resulting in the global changes in gene regulation (Barchuk et al. 2007; Dickman et al. 2013; Ashby et al. 2016). Interestingly, although royal jelly is nutritionally rich with a high proportion of proteins, carbohydrates, fatty acids, and vitamins and minerals (Yu et al. 2023), its effect on honey bees’ gut biochemical environment and microbiota remains unexplored. However, there are studies showing the impact of admission of the royal jelly on mice’s health and their microbiota (Chi et al. 2021). Here, we hypothesize that the overproduction of royal jelly by young nursing bees combined with a low ratio of young larvae in the colony leads to royal jelly cross-feeding among young bees. By utilizing the cutting-edge method for simultaneous microbiota metabarcoding and quantification using amplicon sequencing of two hypervariable regions of bacterial 16S marker gene, we aimed to track changes in young worker bees’ microbiomes throughout swarming preparation.

## Materials and methods

### Specimen collection and preparation

Specimens used in this study came from three experimental colonies in which queens were artificially inseminated by drones originating from a single queen. In early May 2020, after the first drone brood observation, freshly emerged honey bee workers were marked every three days (Suppl. Table 1, Fig. 1). After ten days, worker bees from each batch were collected from the frame, immediately placed on dry ice, and put in -60°C until processed. Next, brains were collected for RNA and ATAC-seq analyses, whereas the rest of the body was stored in 96% ethanol and -20°C for further DNA extraction. Finally, 78 specimens from three colonies were chosen for microbiome analysis (Suppl. Table 1).

**Figure 1.**
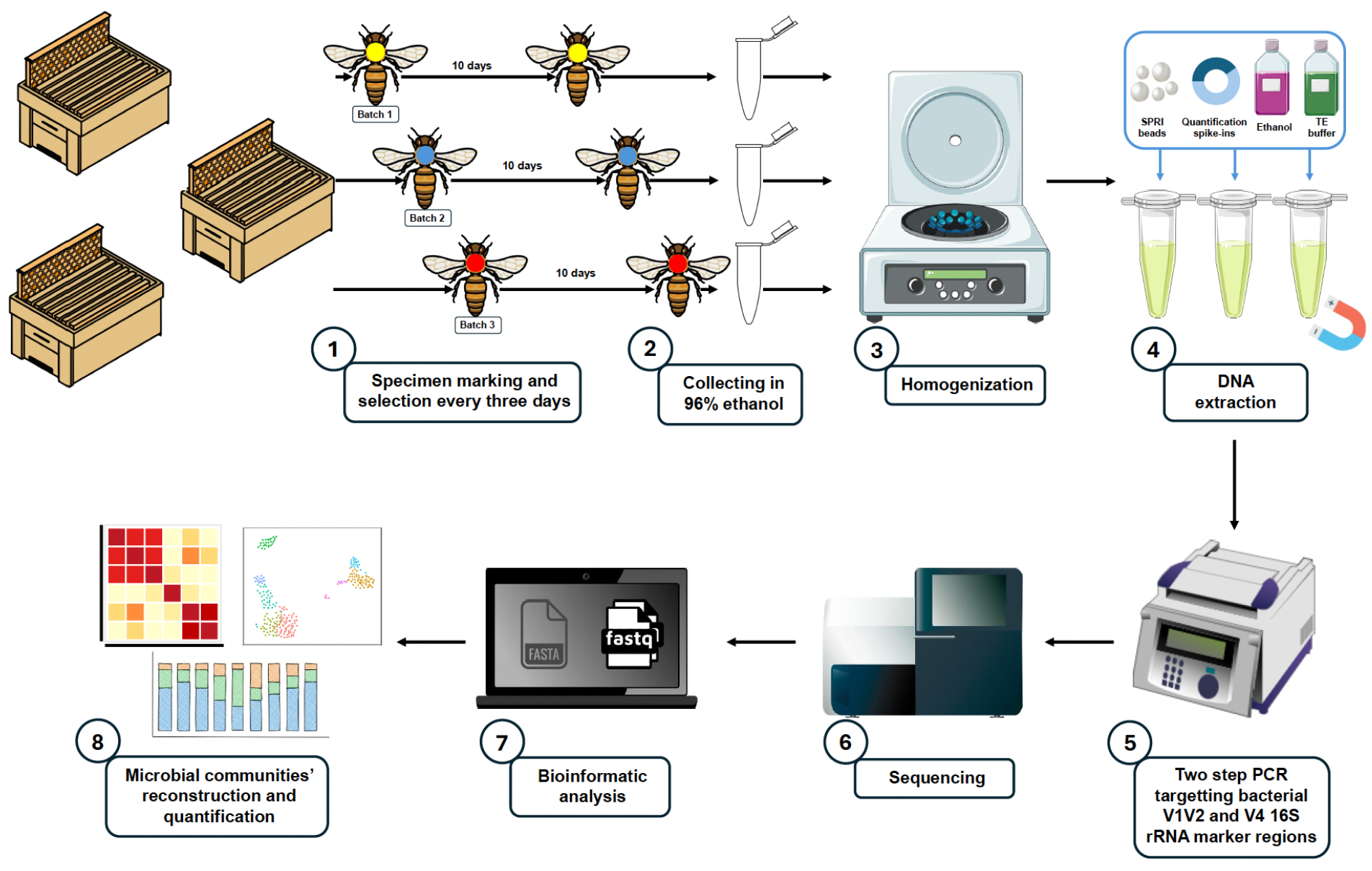
The experimental workflow of the study.

### Lysis and DNA extraction

The worker bees were homogenized by placing them in 2 ml tubes containing 200 µl of a buffer mixture, which included 195 µl of ‘Vesterinen’ lysis buffer (0.4 M NaCl, 10 mM Tris-HCl, 2 mM EDTA pH 8, 2% SDS) (Aljanabi and Martinez 1997; Vesterinen et al. 2016) and 5 µl of proteinase K. Additional buffer was added if necessary to ensure the samples were fully submerged. Ceramic beads (2.8 mm and 0.5 mm) were added to each tube, and the samples were homogenized using the Omni Bead Raptor Elite homogenizer for two 30-second cycles at a speed of 5 m/s. The samples were then incubated at 55°C for 2 hours in a thermal block.

After cooling, 40 µl of the homogenate from each tube was transferred to a deep-well plate. A 100,000 copies of quantification spike-in was added to each sample — a linearised plasmid containing an artificial 16S rRNA target Ec5001 (Tourlousse et al. 2016) in 2 µl of TE buffer. DNA was purified using 80 µl of SPRI beads on a magnetic stand, followed by two washes with 80% ethanol. The DNA was then diluted with 20.5 µl of TE buffer, and 20 µl of this solution was transferred to a new 96-well plate for DNA concentration measurement using the Quant-iT PicoGreen kit.

### Library preparation and sequencing

Amplicon libraries were prepared using a custom two-step PCR protocol (Buczek and Kolasa et al., 2024). In the first step, template-specific primers with Illumina adaptor tails were used: 515F and 806R for the V4 region (Parada et al. 2016) and 27F and 338R for the V1V2 region (Walker et al. 2015) of bacterial 16S rRNA gene. The PCR solution consisted of 5 µl of QIAGEN Multiplex Master Mix, a mix of primers at concentrations 2.5uM (COI) and 10uM (16S V4), 2 µl of DNA template, and 1 µl of water (final volume: 10 µl). The temperature program for the first round of PCR included the initial step of denaturation at 95°C for 15 minutes, followed by 25-27 cycles of denaturation (30s, 94°C), annealing (90s, 50°C) and extension (90s, 72°C) phases, and the final extension step (10m, 72°C). The products were checked on 2.5% agarose gel against positive and negative controls and cleaned with SPRI beads.

Illumina adapters and unique index pairs were added to the samples during the second indexing PCR. The temperature program for PCR in this step remained the same, but the number of cycles was reduced to 7. In each laboratory step (DNA extraction, first and indexing PCR), we added a negative control (blanks).

The libraries were pooled approximately equimolarly based on band intensity on agarose gels to ensure a roughly equal representation of each sample in the pool. After the last cleaning step with SPRI beads, the pools were ready for sequencing performed on an Illumina MiSeq v3 lane (2 × 250 bp reads) at the Institute of Environmental Sciences of Jagiellonian University.

### Bioinformatic analysis

The obtained data was analyzed by a set of custom Python scripts developed in the Symbiosis Evolution Research Group at Jagiellonian University (Buczek et al. 2024). All scripts that were used in this project are available at the GitHub repository: https://github.com/MikeCollasa/Swarming_project.

First, R1 and R2 files for each library were split into bins corresponding to the target marker region, and primers were cut off. Due to the unique set of informative indexes used for each sample, the potential cross-talk (Wright and Vetsigian 2016) between libraries was removed, leaving only the reads characteristic of a particular sample. Next, R1 and R2 reads were joined into high-quality contigs (minimum Phred score 30) using PEAR (Zhang et al. 2014). Subsequently, contigs were de-replicated (Rognes et al. 2016) and denoised (Edgar 2016) separately for each of the libraries to avoid losing biologically relevant information about rare genotypes that could happen during the denoising of the whole dataset at once (Prodan et al. 2020). Chimeras were recognized using USEARCH, and each sequence was taxonomically classified using the SINTAX algorithm and customized SILVA database (version 138 SSU) (Quast et al. 2013). Afterwards, sequences were grouped based on a 97% similarity threshold into Operational Taxonomic Units (OTUs) with the UPARSE-OTU algorithm implemented in USEARCH. At this stage of the analysis, two tables were produced: ASVs table (Amplicon Sequencing Variant) (also known as zOTUs - zero-radius Operational Taxonomic Units) describing genotypic diversity and OTUs table (Operational Taxonomic Units) – clustering genotypes based on an abovementioned similarity threshold. Using negative controls generated in each laboratory step, bacterial 16S rRNA gene data were screened for putative DNA extraction and PCR reagent contaminants. We first removed genotypes classified as chloroplasts, mitochondria, Archea, or chimers using information about taxonomy classification. Next, we calculated relative abundances and used ratios of each genotype presented in blank and experimental libraries to accurately assign genotypes as putative real insect-associated microbes or PCR or extraction contaminants.

Next, reads identified as quantitative spike-ins were used to reconstruct bacterial absolute abundances in the processed honey bee workers. Specifically, the symbiont-to-extraction spike-in ratio, multiplied by the number of extraction spike-in copies and the proportion of the homogenate, allowed us to estimate amplifiable bacterial 16S rRNA copy numbers in the homogenized specimens (Buczek et al. 2024).

Finally, manual analysis was conducted to remove controls and samples with incorrect indexes or zero abundance of bacteria and create the dataset used in the statistical analysis.

### Statistical analysis and visualization

Statistical analysis was performed using RStudio version 2023.03.1+446 (R Core Team, 2023), and Processing 3 software version 3.5.4 (Reas and Fry 2006) was used for generating heatmap. Inkscape 1.2.2 (Inkscape Project, 2022) was used to modify generated plots and visualizations. All pictures used for the methodological scheme (Fig. 1) come from https://bioicons.com/website.

To evaluate the impact of batch effects on the absolute abundances of microbial communities, we used a linear mixed effects model (LME) with log-transformed absolute abundance as a response variable, batch as a fixed effect, and hive as a random effect. The model was fitted using restricted maximum likelihood (REML) estimation. The analysis used the lme4 package (Bates et al. 2015). The model’s fit was assessed through the REML criterion at convergence, and the significance of fixed effects was evaluated using Satterthwaite’s method for degrees of freedom, as implemented in the lmerTest package (Kuznetsova et al. 2017).

The Bray-Curtis (Sørensen 1948) dissimilarity matrix was computed from the absolute abundance data. This distance matrix quantifies the compositional dissimilarity between pairs of samples based on their zOTU profiles. To assess the influence of temporal changes (represented by different batches) on microbial community composition, a PERMANOVA (Anderson 2001) was performed using the adonis function from the vegan (Oksanen et al. 2024) R package. We implemented differential abundance analysis to verify the changes of particular bacterial taxa absolute abundance throughout swarming preparation, which was performed using the DESeq2 R package (Love et al. 2014). The DESeq2 package was used to fit a negative binomial model to the count data and to perform Wald tests (Gourieroux et al. 1982) for differential abundances between each batch.

A Bland-Altman plot (Dewitte et al. 2002) was generated using the ggplot2 R package (Wickham 2016) to assess the agreement between two quantification methods (based on two gene fragments: 16SV1V2 and 16SV4). The mean and difference between the log-transformed values of the two methods were calculated for each sample. The mean difference, known as bias, was computed, along with the 95% limits of agreement, defined as the mean difference ± 1.96 times the standard deviation of the differences.

## Results

### Data processing and decontamination

#### 16S rRNA V4 dataset

After bioinformatic analysis, two samples were excluded due to insufficient read number and quality, and 76 experimental libraries remained. The number of reads before decontamination ranged between 24,286 and 183,537 (Suppl. Table 2.1). After the decontamination, the number of reads ranged between 161,303 and 23,788, providing sufficient depth for further analysis (Suppl. Table 2.2). The two most abundant non-bacteria zOTUs (zOTU7, OTU6, and zOTU14, OTU9) were classified by our pipeline as chloroplast and mitochondria, respectively. After further blasting, the top hits indicated that zOTU7 was classified as belonging to the family Rosaceae (GenBank accession number: MW755928.1), whereas zOTU14 was *Malus domestica* (GenBank accession number: OX352782.1). Both genotypes accounted for 6.3% of all reads across all experimental libraries. zOTU7 had 261,610 reads across all experimental samples, with relative abundance ranging between 14.95%% and 0.05%, whereas zOTU14 had 133,279 reads across all experimental samples, with relative abundance ranging between 7.00% and 0.03% (Suppl. Table 2.3 and 1.6). Genotypes assigned as contaminants or non-bacteria were represented by 573,059 reads across all libraries, constituting 9.1% of the dataset (Suppl. Table 2.3 and 1.6). The most abundant reagent contaminant was zOTU26 (OTU14), which was taxonomically delimited as *Brachybacterium*. 32,612 reads represented this genotype and were responsible for 0.5% of all reads across libraries (min 0%, max 0.36% in libraries) (Suppl. Table 2.1).

#### 16S rRNA V1-V2 dataset

After bioinformatic analysis, similarly to the 16SV4 dataset, two samples were excluded due to insufficient read number and quality, and 76 experimental libraries remained. The number of reads before decontamination ranged between 8505 and 98,870 (Suppl. Table X). After the decontamination, the number of reads ranged between 95,424 and 7986. Although the V1V2 dataset had lower read numbers than the V4 dataset, the numbers were still sufficient for further analysis (Suppl. Table X).

The three most abundant zOTUs assigned as non-bacteria (zOTU5, zOTU55 and zOTU112) were members of the same OTU4 and assigned as chloroplast. However, further blasting didn’t provide congruent results as those different zOTUs were classified as various plant species. Those three genotypes were responsible for 3.98% (max 8848 reads in a library), 0.29% (max 1341 reads) and 0.09% (max 318 reads), respectively, of the whole dataset abundance (Suppl. Table 3.1). Overall, zOTUs assigned as non-bacteria had 167,779 reads and constituted 5.81% of the whole dataset (Suppl. Table 3.1), and all genotypes assigned as contaminants were present only in blank samples.

### Microbial communities’ composition in same-age worker bees throughout swarming preparation

#### 16S rRNA V4 dataset

After decontamination, we identified 26 OTUs that were assigned by our decontamination pipeline as symbionts (Suppl. Table 2.4, Fig. 2). Six out of those were low abundant OTUs (<0.001 of maximum relative abundance across all libraries) and taxonomically assigned as *Rahnella, Pseudomonas* (two OTUs) (Suppl. Table 2.5), *Serratia, Sphingomonas*, and *Methylobacterium*. 17 OTUs reached a threshold of at least 1% relative abundance across all libraries or belonged to a known core microbiome taxa of honey bees (Suppl. Table 2.5, Fig. 2). Regarding genotype diversity, we reconstructed 205 zOTUs of the 26 OTUs above. 69 out of all zOTUs (belonging to 12 OTUs) were responsible for at least 1% of relative abundance in any of the samples. In the context of taxa belonging to honey bees’ core microbiota, the most diverse were symbionts assigned as *Lactobacillus* with seven OTUs and 103 zOTUs in total (39 of which were present in at least one sample with relative abundance >1%). *Bifidobacterium* represented by a single OTU had 23 unique variants, with eight reaching at least a 1% threshold in any of the libraries. *Snogdrasella* was represented by 17 genotypes belonging to a single OTU, with 7 representing at least 1% relative abundance across all libraries. Similarly, *Gilliamella* constituted a single OTU with ten zOTUs (six with relative abundance >1%). Additionally, we found OTUs previously described as honey bees’ symbionts. *Frischella* was represented by a single OTU with ten zOTUs, out of which five had at least 1% relative abundance among samples. We found a single *Acetobacteraceae* OTU with five unique genotypes, with two passing a 1% threshold. *Bombella* and *Fructobacillus* were also represented by a single OTU (with two and three zOTUs, respectively); however, their abundance didn’t reach the 1% threshold (Suppl. Table 2.3).

**Figure 2.**
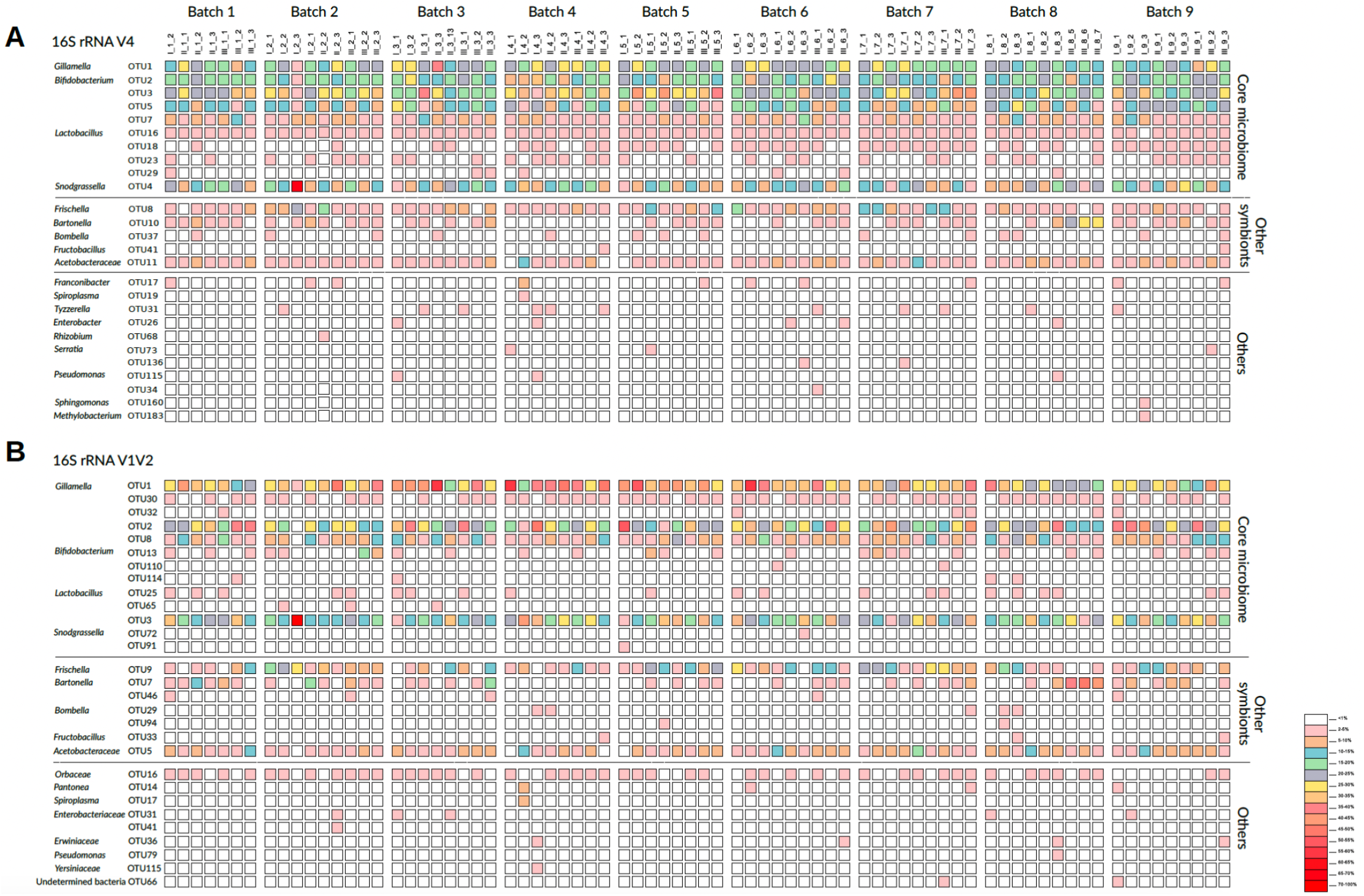
The bacterial relative abundance across studied batches. A. Reconstruction based on bacterial 16S rRNA hypervariable region V4. B. Reconstruction based on bacterial 16S rRNA hypervariable region V1V2.

#### 16S rRNA V1-V2 dataset

In the decontaminated dataset, we identified 29 OTUs, of which 15 exceeded at least 1% relative abundance in any of the experimental libraries (Fig. 2, Suppl. Table 3.4 and 2.5). The known members of honey bee microbiota consisted of *Snodgrassella*: OTU3, OTU72, OTU91 (maximum relative abundance in any of the library: 70.73%, 0.24%, 0.1% respectively); *Lactobacillus*: OTU25, OTU65 (max abundance: 0.4%, 0.24%); *Bifidobacterium*: OTU2, OTU8, OTU13, OTU110, OTU114 (max abundance: 50.26%, 23.85%, 15.61%, 0.19%, 0.07%); *Gilliamella*: OTU1, OTU30, OTU32 (max abundance: 55.56%, 3.20%, 0.34%); *Fructobacillus* OTU33 (max abundance 1.71%; Acetobacteraceae OTU5 (max abundance: 19.77%); *Bartonella*: OTU7 and OTU46 (max abundance: 49.14%, 0.41%) and *Bombella* OTU29 and OTU94 (max abundance: 0.62%, 0.13%), *Frischella* OTU9 (max abundance: 34.92%). Other symbionts belonging to *Pantoea* (OTU14), *Spiroplasma* (OTU17) - classified as *Spiroplasma melliferum* (top hit GenBank accession number: CP029202.1), Enterobacteriaceae (OTU31 and OTU41), Orbaceae (OTU16), Erwiniaceae (OTU36), undetermined bacteria (OTU66), Yersiniaceae (OTU115), *Pseudomonas* (OTU 79) represent a maximum abundance in any of the libraries ranging from 0.06% (*Pseudomonas*) to 9.38% (*Pantonea*) (Fig. 2, Suppl. Table 3.5). The V1V2 dataset was more abundant in terms of genotypic diversity than the V4 dataset, with 869 unique genotypes and 208 (describing at least 1% relative abundance and belonging to 14 OTUs). *Snodgrassella* was described by 65 zOTUs (63 belonged to OTU3, and 19 of them exceeded 1%); *Gilliamella* consisted of 166 zOTUs (67 with at least 1% abundance); *Lactobacillus* OTU25 had nine zOTUs and OTU65 three, none of which exceeded 1%; *Bifidobacterium* with 265 zOTUs in total (OTU2: 129 genotypes, 29 with >1% abundance; OTU8: 99 genotypes, 26 with >1%; OTU13: three with >1% abundance; OTU 110 had two and OTU114 had six genotypes and with none exceeding 1% threshold; Acetobacteraceae OTU5 consisted of 22 zOTUs (four with maximum abundance in any of the samples>1%); *Bartonella* OTU7 had 52 genotypes (14 with >1% abundance) and OTU46 consisted of seven zOTUs none of which didn’t pass 1% threshold; *Bombella* OTU29 and OTU94 were both represented by two genotypes with maximum relative abundance less than 1% and *Frischella* OTU9 with 120 zOTUs (36 with >1% abundance)(Suppl. Table 3.2 and 2.3).

### The impact of the batch effect on microbial quantities

#### 16S rRNA V4 dataset

The absolute abundance (presented as log_10_) ranged between 7.06 and 9.19, with a mean value of 8.22 (Suppl. Table 2.7, Fig. 3). In our linear mixed model analysis of absolute abundances, we found no significant batch effects (p = 0.735), indicating that temporal variations between batches did not substantially impact the measured absolute abundances of microbial communities (Suppl. Table 2.8). Hence, we checked if batch factor can influence microbial composition at both OTU and zOTU levels. We conducted a PERMANOVA to evaluate the effect of batch on absolute microbial community composition. The analysis revealed a significant impact of the batch group on the community structure for both OTU (F = 1.672, R^2^ e= 0.1664, p = 0.025) and zOTU (F = 1.5112, R^2^ = 0.1529, p = 0.013) datasets (Suppl. Table 2.9). The differential abundance analysis identified ten OTUs with significant changes between batches. However, only two (OTU17 *Franconibacter* and OTU31 *Tyzzerella*) changed in at least four consecutive batches (Suppl. Table 2.11). On the other hand, zOTU16 (OTU10, *Bombella*) was the only genotype with significant differences across multiple batch comparisons (Suppl. Table 2.10). However, its abundance changed consecutively only between batches three to five (Suppl. Table 2.10).

**Figure 3.**
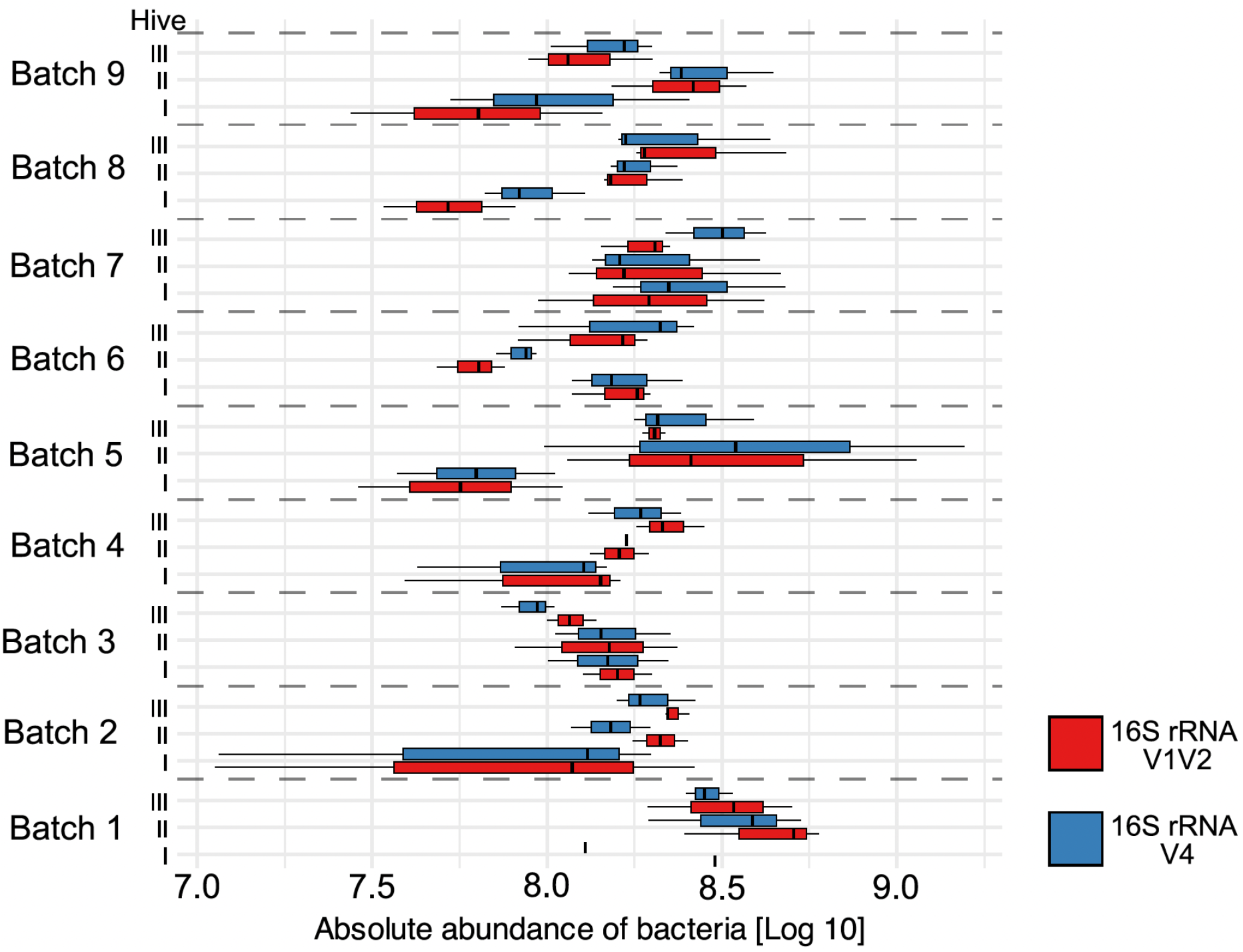
Comparison of the absolute abundance of bacteria between batches and used marker gene regions.

#### 16S rRNA V1-V2 dataset

Due to the lack of spike-in reads in one of the samples (II_2_2), it was excluded from absolute abundance analysis. The absolute abundance (presented as log_10_) ranged between 7,05 and 9,05, with a mean value of 8,19 (Suppl. Table 3.7, Fig. 3). The V1V2 provided congruent results as the V4 dataset, as the linear mixed model analysis showed no significant batch effect (p =0.281) on absolute abundance between batches (Suppl. Table 3.8). Similarly, the PERMANOVA analysis revealed a significant impact of the batch group on the community structure for both OTU (F = 1.8411, R^2^ = 0.1824, p = 0.006) and zOTU (F = 1.2528, R^2^ = 0.1318, p = 0.009) datasets (Suppl. Table 3.9). Interestingly, differential abundance analysis conducted on the V1V2 dataset identified six OTUs with significant changes across batches, but only one (OTU29 *Bombella*) changed across at least four consecutive batches (Suppl. Table 3.11). On the other hand, we found 193 zOTUs with significant abundance changes across batches. However, only 23 had significant changes across more than four consecutive batches: *Gilliamella* OTU1 (seven zOTUs with changes in five to seven batches), OTU16 and OTU30 (both having one genotype with six changes), *Bifidobacterium* OTU2 (three genotypes with changes in five batches), OTU8 (five unique sequences with five to seven changes) and OTU13 (one zOTU with five changes), OTU3 *Snodgrassella* and OTU7 *Bartonella* (both with one zOTU with five changes), OTU9 *Frischella* (three genotypes with five and seven changes). The number of reads for mentioned zOTUs ranged between 233 and 35,712 across the whole dataset, and their maximum abundances in any of the libraries varied between 0.19% and 34.07% (Suppl. Table 3.10).

#### Quantification method comparison between datasets

The Bland-Altman plot (Suppl. Fig. 1) was generated to assess the agreement between the two quantification methods, based on 16SV12 and 16SV4, after the log transformation of the data. The mean difference (bias) between the log-transformed values was approximately 7.5×10^9^, indicating a notable positive bias, where the V12 quantification tends to yield higher values than the V4 method. The 95% limits of agreement were calculated as approximately 2.5×10^9^ (lower limit) and 1.25×10^9^ (upper limit). Most of the differences between the two methods fell within these limits, suggesting reasonable agreement across the range of values, though some variability was observed, particularly at higher mean values. This trend suggests that while the two methods largely agree for smaller measurements, discrepancies arise as the values increase.

## Discussion

### Challenges in amplicon-based microbiome studies

In recent years, microbial studies have gained much popularity due to lowered library preparation and sequencing costs and the availability of user-friendly tools for easy amplicon data analysis. Although metagenomic approaches are becoming more available in terms of price, they still present challenges due to their complex and multi-aspect analysis. Hence, the preliminary studies based on amplicon data followed by a more precise analysis using metagenomic approaches have been proposed as the most efficient way to study the dynamics between hosts and their microbial allies (Łukasik and Kolasa 2024; Buczek et al. 2024). Nonetheless, amplicon data analysis contains multiple methodological caveats (Knight et al. 2018) that must be addressed appropriately to draw proper biological conclusions. However, most of the studies still suffer from such issues. In fact, a recent metanalysis (Williamson et al. 2024) pointed out that more than 86% of studies on insect microbiomes conducted between 2001 and 2022 might provide biased information about the microbiome composition and its potential function on insect hosts due to lack of careful decontamination. Therefore, in our study, we implemented rigorous laboratory steps (multiple blank samples on each laboratory step) and bioinformatic procedures (denoising every library separately and implementing a custom decontamination tool) to efficiently filter out contaminants and PCR artefacts that may introduce false biodiversity into the datasets.

### Lack of changes in microbial quantities throughout swarming preparation

Insect species differ dramatically in the abundance of microorganisms they host, and these differences often correlate with the microbes’ function in insect biology (Hammer, Sanders & Fierer, 2019). In the case of healthy honey bees, the estimates suggest a stable and constant abundance on a level between 10^8^ and 10^9^ (Motta and Moran 2024). However, most studies fail to report bacterial absolute abundance with a few notable exceptions (e.g., Schwarz et al. 2016; Kešnerová et al. 2020; Castelli et al. 2022), leaving a gap in understanding the quantitative dynamics of bee-associated microbiota.

Here, we used a new approach to quantify the microbial community as additional information without increasing the costs of multitarget amplicon sequencing (Buczek et al. 2024), which opens up new possibilities to study insect-microbiome interactions on a new higher level. The minimum and maximum absolute abundance estimates were congruent between the V1V2 and the V4 datasets, ranging between 10^7^ and 10^9^. This consistency supports the accuracy of our method, and it also shifts the described abundance (Motta and Moran 2024) to lower values. The lack of changes in overall microbiome abundance with simultaneous changes in microbial composition might suggest that physiological factors allow honeybees to sustain bacterial quantities on a particular stable level during the swarming preparation. We can not also exclude the possibility that the stability in microbiome quantities among the analyzed bees is related to their age. Specifically, since we used 10-day-old worker bees, the overall homogeneity of the abundances might be the outcome of the relatively short time from their emergence from pupae as the adult honey bees emerge sterile and acquire symbionts through trophallaxis with other nestmates (Moran 2015). As the detected minimal absolute abundance (10^7^) was one order of magnitude lower than previously reported (10^8^), it would be possible that the quantities of relatively young bee workers do not reach their optimal abundances yet and may increase in the following days of their life. On the other hand, Kešnerová et al. (2020) demonstrated the seasonal variability in bee microbiota abundance, revealing statistically significant differences in total microbiome load between winter bees and foragers. Notably, our results correspond well with their findings, showing that foragers sampled in May and June (the same month we sampled our bees) tend to have lower abundancies (10^7^). Thus, it is possible that the microbial quantities may be influenced not only by the bee’s age (behavioural tasks) but also by season-dependent environmental factors such as pollen source and quality.

### Overall microbiome composition in same-age nursing bees throughout swarming preparation

The microbial community of honey bees is relatively consistent in terms of composition; however, beyond the core microbiota, these insects may host additional bacteria whose presence may be host-specific and related to various factors. In this study, we analyzed the honey bee microbiome using the V1V2 and V4 regions of the bacterial 16S rRNA gene, revealing slight differences in microbiome composition depending on the region used. Our results confirmed previous findings showing that relying on a single hypervariable region of 16S rRNA is not sufficient for accurate reconstruction of honey bee microbiome with the V1V2 region complementing the widely used V4 region (Romero et al. 2019). Nonetheless, both datasets consistently identified five core clusters characteristic of the microbiome of honey bees: *Snogdrassella, Bifidobacterium, Lactobacillus* (currently distinguished into *Lactobacillus* and *Bombilactobacillus*), and *Gillamella* (Motta and Moran 2024; Luo et al. 2024). Additionally, we captured honey bees’ symbionts that are not included in the core microbiome: *Frischella, Bartonella, Bombella, Fructobacillus*, and bacterium belonging to the Acetobacteracae group (Motta and Moran 2024). However, we observed a clear bias in terms of OTU diversity. The V4 primers distinguished more OTUs within the *Lactobacillus* group, whereas V1V2 primers were better at detecting *Snodgdrasella, Gillamella, Bifidobacterium, Bartonella*, and *Bombella*. Surprisingly, we did not find *Commensalibacter*, the taxon typically present in honey bee microbiota (Motta and Moran 2024). However, this might be explained by results indicating that this taxa is much more prevalent in winter bees than spring/summer bees (Kešnerová et al., 2020). Other bacteria often described as parts of the honey bee microbiome, including *Pantonea, Serratia* (only in the V4 dataset), and Enterobacteriaceae (Kešnerová et al. 2020; Motta and Moran 2024) were also found in our datasets. Interestingly, we found *Spiroplasma* sequences prevalent in one of the samples. Although blasting it against the NCBI database indicated it as *Spiroplasma melliferum*, we interpret it more as a possible signal from one of the arthropod parasites such as *Varroa destructor* or tracheal mite (*Acarapis woodi*). 16S rRNA signal from endosymbionts colonizing parasites or parasitoids has been previously described as factors blurring the true microbiome composition of insects (Kolasa et al. 2023).

### Changes in microbial composition throughout swarming preparation

Previous studies have shown that microbiome changes in honey bees are not dependent on the location of the apiary but rather on the time point at which bees were collected (Almeida et al. 2023). Our results support that claim, showing a significant effect of the batch on microbial composition with no effect of the hive factor. Results of our analyses indicate that depending on the dataset (V1V2 vs V4) and clustering level (zOTUs vs OTU), different bacterial taxa or genotypes can be distinguished with their absolute abundance shifting through the swarming preparation. In the V4 dataset, we identified two OTUs with statistically relevant changes: *Franconibacter* and *Tyzzerella*. The first one was described as a member of a saccharification agent used to initiate fermentation in the production of Chinese liquor and vinegar (Gao et al. 2017), whereas the second was recently proposed as a factor playing a potential role in larvae biology (Maigoro et al. 2024). Hence, although they might be members of honey bees’ microbiota, considering their low abundance and the limitations of amplicon-based bacterial metabarcoding, we would be far from assessing their significant role in triggering the swarming preparation or playing an essential role in the process. On the other hand, the changes in the prevalence of *Bombella* zOTU might indicate its function in this process. *Bombella* has been recognized as a diverse taxon with different strains contributing to various aspects of honey bees’ larval and adult biology, ranging from protection against fungal pathogens to larvae supplementation in lysine and buffering nutritional stress (Parish et al. 2022; Miller 2023). Lysine is directly involved in nitric oxide synthesis, a neurotransmitter affecting honey bees’ brain functions (Gage et al. 2020).

The V1V2 dataset analysis showed much more variation on both zOTU and OTU levels. In the zOTU dataset, we found 23 genotypes with significant changes in at least four consecutive batches. Those genotypes belonged to *Gilliamella, Bifidobacterium, Snodgrassella*, and *Bartonella* groups. Considering that the V4 dataset showed a stable diversity pattern of the first three species (belonging to the core microbiome), we tend to interpret those changes as an outcome of a bias caused by V1V2 primers to overrepresent the genetic diversity of those taxa. *Bartonella* was a bacterial symbiont captured in the V1V2 dataset but absent in the V4 dataset. This bacteria has been described as especially abundant in workers involved in hive tasks such as food processing (Jones et al. 2018b) and requiring food rich in pollen to grow in abundance successfully (Kešnerová et al. 2020). Its change over the sampling period might be explained by a higher pollen supply available from the environment between May and June. The genomic-based reconstruction of *Bartonella* metabolism showed that it is capable of tryptophan and phenylalanine secretion (Li et al. 2022). Tryptophan regulation by *Lactobacillus* has been shown to influence the memory behavior and learning of worker bees (Zhang et al. 2022a), suggesting its potential role in triggering the changes in behavior during swarming preparations. Interestingly, the only zOTU showing statistically significant changes in at least four consecutive batches was *Bombella*, which overlaps with results obtained by analyzing the OTU V4 dataset.

## Conclusions

Honey bees’ taxonomically stable microbiota, with its potential strain diversity combined with their complex behavior, makes them the perfect model species for studying gut-brain interactions. We tested the hypothesis that the gut-brain axis can trigger swarming preparation by implementing a novel approach for simultaneous microbiota metabarcoding and its quantification based on multitarget amplicon sequencing. Our results indicate that it is plausible that some taxa may play a role in changing the behavior of young honey bees, although further studies involving gene expression analysis and genome reconstruction of a symbiont strain would be required, similar to an approach used in ants showing the correlation between microbiome composition and brain gene expression (Kay et al. 2023).

## Data availability

All data underlying this study have been deposited in the Sequence Read Archive (SRA) of the National Center for Biotechnology Information (NCBI) under BioProject Accession no. PRJNA1161854.

## Acknowledgements

We thank Monika Prus-Frankowska for her support in implementing laboratory procedures and Pasieka Szeligów for providing inseminated queens. The Polish National Science Centre grant 2018/31/N/NZ8/03406 supported the project.

## Statements

The authors have no relevant financial or non-financial interests to disclose.

