## Supplementary figures and images for "Peasants at the queen’s table?: The microbiome’s dynamics throughout swarming preparation in honey bees (*Apis mellifera*)"

### Suppl. Fig. 1

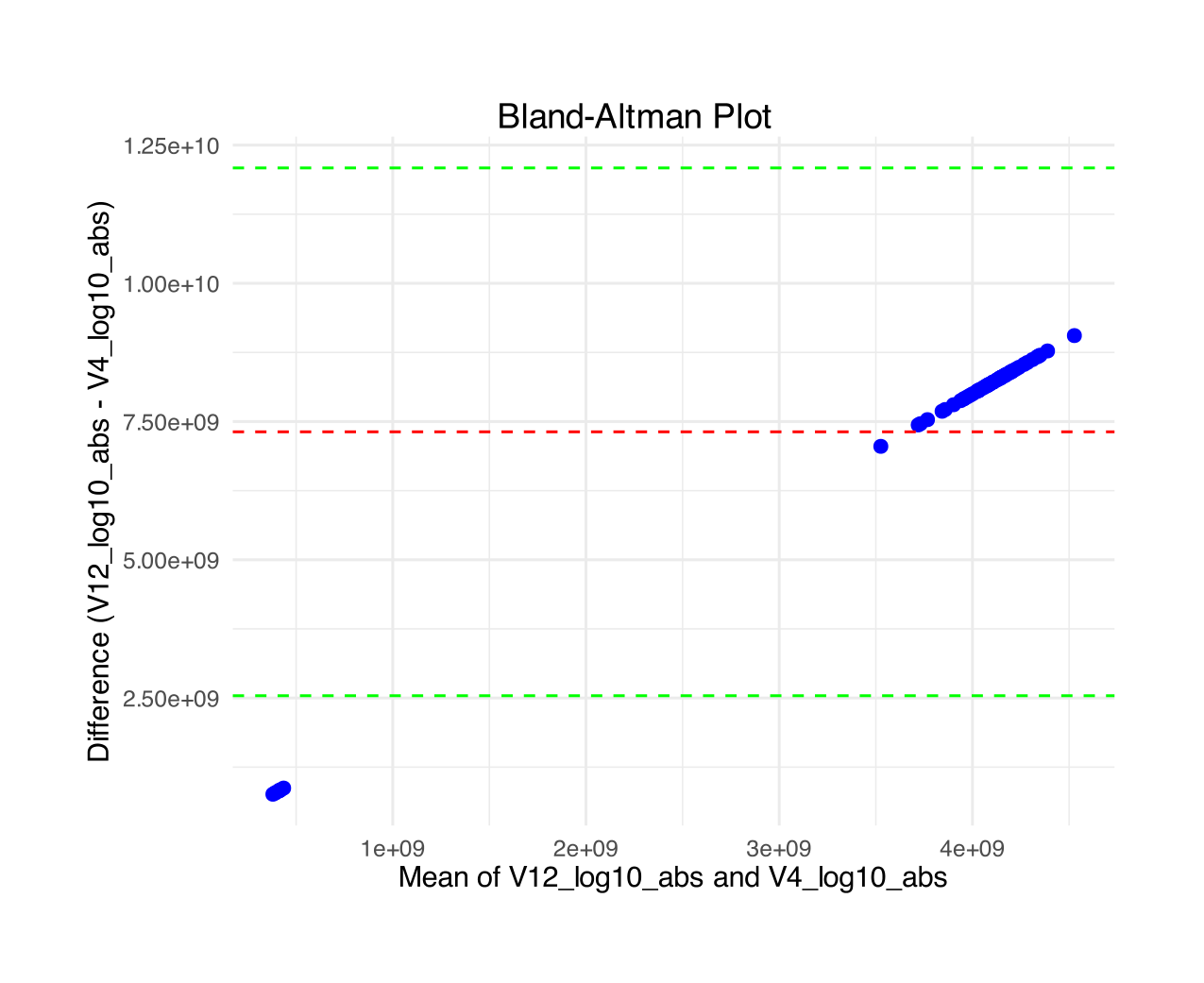
